# *Igf2* regulates early postnatal DPP4^+^ preadipocyte pool expansion

**DOI:** 10.1101/2025.02.08.637214

**Authors:** Irem Altun, Safal Walia, Xiaocheng Yan, Inderjeet Singh, Ruth Karlina, Viktorian Miok, Lingru Kang, Khanh Ho Diep Vo, Dominik Lutter, Fabiana Perrochi, Siegfried Ussar

**Affiliations:** RU Adipocytes & Metabolism, Helmholtz Diabetes Center, Helmholtz Zentrum München, German Research Center for Environmental Health GmbH, 85764 Neuherberg, Germany; German Center for Diabetes Research (DZD), 85764 Neuherberg, Germany; Institute for Diabetes and Obesity, Helmholtz Diabetes Center, Helmholtz Zentrum München, German Research Center for Environmental Health GmbH, 85764 Neuherberg, Germany; Munich Cluster for Systems Neurology, Munich, Germany; Institute of Neuronal Cell Biology, Technical University of Munich, Munich, Germany

**Keywords:** adipose tissue development, preadipocyte, IGF2, DPP4+ Progenitor Cells, Hyperplasia

## Abstract

Adipose tissue is rapidly expanding early in life. Identifying the queues facilitating this process will advance our understanding of metabolically healthy obesity. Single-cell RNA sequencing revealed compositional differences between pre-wean and adult murine subcutaneous adipose tissue. We identify a dipeptidyl peptidase-4 (*Dpp4*) positive precursor population in pre-wean mice residing at the reticular interstitium of subcutaneous adipose tissue expressing insulin growth factor 2 (*Igf2)*. We show that IGF2 drives proliferation rather than differentiation in these cells. Moreover, loss of *Igf2* in *Dpp4*^+^ progenitor cells promotes adipogenesis. We show that expression of *Igf2* at a limited timeframe during development promotes preadipocyte expansion.

## Introduction

White adipose tissue (WAT) development begins in utero and continues after birth (Wang et al. 2013). In humans a rapid and massive expansion of adipose tissue, mainly by the increase in adipocyte size, peaks within the first year of life followed by a significant decrease after two years postpartum (Knittle et al. 1979). In line with this, murine subcutaneous adipose tissue mass, relative to body weight, peaks at 12 days postpartum and decreases proportionally during weaning (Tsukada et al. 2023). Differences in adipocyte size and composition before, during and after weaning further support developmental changes and remodeling of the tissue during early life (Zhang et al. 2022; Tsukada et al. 2023; Qian et al. 2024). This reconfiguration is critical as the continued expansion of adipose tissue into adolescence results in obesity in adulthood (Knittle et al. 1979). However, the regulation of this initial, metabolically healthy, adipose tissue expansion early in life remains incompletely understood (Altun et al. 2022).

WAT is a heterogenous endocrine organ. It consists of immune, adipose progenitor (APC), endothelial and neuronal cell populations (Suwandhi et al. 2021; Altun et al. 2022; Emont et al. 2022). Several studies identified distinct APC subtypes with different functions in adipose tissue physiology. Sakers *et al*. defined APCs as highly proliferative cells with stem-like properties that maintain the progenitor niche in the tissue (Altun et al. 2022; Sakers et al. 2022). Merrick *et al*. identified a DPP4^+^ progenitor cell population that gives rise to committed preadipocytes (Merrick et al. 2019). These cells, residing in the reticular interstitium (RI), an area surrounding adipose tissue, have an increased proliferation rate. In addition, some precursors directly promote adipogenesis (Rodeheffer et al. 2008; Merrick et al. 2019; Altun et al. 2022). However, what regulates the proliferative capacity of these cells and induces commitment to differentiation needs further investigation.

Insulin growth factor 2 (IGF2) is a secreted protein that is essential for fetal growth (DeChiara et al. 1991; Sandovici et al. 2022). IGF2 is a member of the insulin superfamily and can activate both insulin receptor (InsR) and IGF1 receptor (IGF1R) signaling pathways (Frasca et al. 1999; Blyth et al. 2022). Both pathways have been described as regulators of adipose tissue development and metabolism (Bluher et al. 2002; Boucher et al. 2012). Even though IGF2 supplementation was shown to regulate adipogenesis in a depot dependent manner, whether APCs are the target and what is the direct function of *Igf2* is not well understood (Alfares et al. 2018).

Here, we studied compositional differences of subcutaneous white adipose tissue (scWAT) between the pre-wean and adult state in mice to discover cell populations or factors regulating healthy adipose tissue expansion. We identified a subpopulation of progenitor cells that are *Dpp4*^+^*Igf2*^+^ residing at the RI in pre-wean mice. DPP4^+^ preadipocytes loose *Igf2* expression later in life together with their high proliferative capacity. Furthermore, loss of *Igf2* function induces differentiation of the DPP4^+^ preadipocytes. Thus, our results propose a mechanism for the regulation and maintenance of progenitor cell expansion.

## Results and Discussion

### *Igf2* is differentially expressed between pre-wean and adult scWAT preadipocytes

We previously reported single cell RNA sequencing (scRNAseq) data comparing pre-wean and adult scWAT stromal-vascular cells (SVCs) (Suwandhi et al. 2021). This analysis revealed a distinct separation of *Pdgfra^+^* preadipocytes between the two age groups (**Fig. S1a**) (Suwandhi et al. 2021). We further analyzed these data to identify new subsets of preadipocytes or individual genes contributing to adipose tissue expansion via hyperplasia. Our analysis revealed 19 cell clusters consisting of a variety of cell types, including immune cells and endothelial cells (**Fig. S1b**). Clusters 1, 2, 5 and 8 expressed elevated levels of preadipocytes markers (*Pdgfra*, *Cd34* and *Dlk1*) (**Fig. S1c**), which were selected for further analysis (**Fig. 1a**). Differential gene expression analysis of pre-wean and adult scWAT preadipocytes revealed *Igf2* as the most differentially expressed gene in pre-wean compared to adult preadipocytes (**Fig 1b,c, Fig S1d**). Further analysis confirmed that *Igf2* mRNA levels were significantly lower in scWAT preadipocytes of adult mice compared to pre-wean mice (**Fig. 1d-g**). The much higher expression of *Igf2* in SVCs compared to the whole tissue, further suggests a primary expression of *Igf2* in preadipocytes. We then assessed the IGF2 signaling pathway in preadipocytes by stimulating cultured primary preadipocytes with 10 nM IGF2. This induced AKT and ERK phosphorylation after 10 minutes compared to controls (**Fig S1e**). The stimulation of these pathways was mediated by activation of both IGF1R and InsR signaling, as shown by phosphorylation of receptor tyrosine residues after IGF2 stimulation (**Fig. S1g-f**).

**Figure 1:**
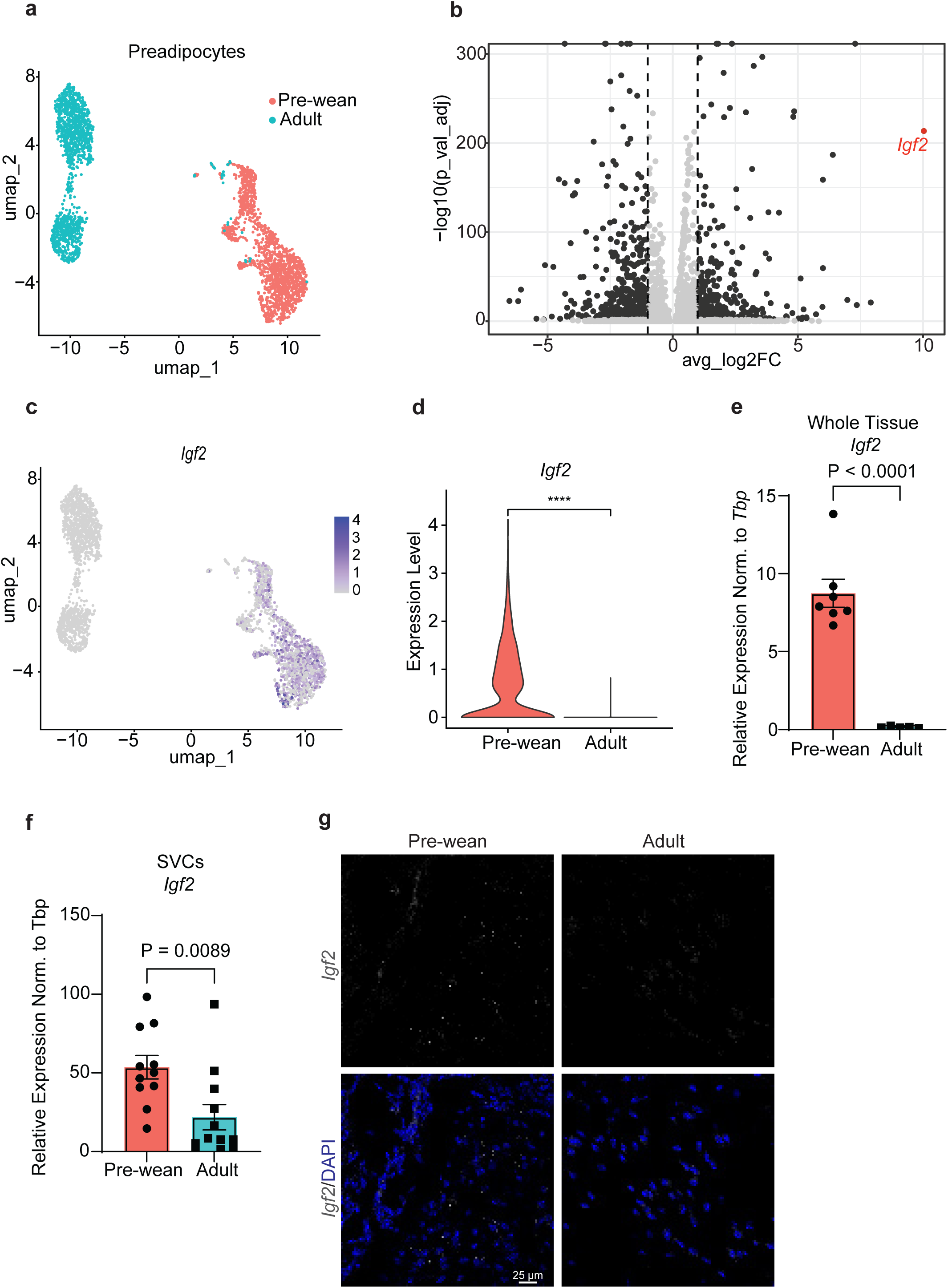
scRNAseq analysis reveals *Igf2* as the most abundantly expressed gene in pre-wean scWAT preadipocytes. (**a**) Projection of two age groups (pre-wean and adult) on scWAT preadipocytes shown as UMAP plot. (**b**) Differential gene expression comparing pre-wean vs adult scWAT preadipocytes. Red dot represents *Igf2*. (**c**) UMAP plot showing the projection of *Igf2* on scWAT preadipocytes. Feature plot colour scales show log-normalized scaled expression values. (**d**) Violin plot showing *Igf2* expression in two age groups. (**e**) RT-qPCR analysis of *Igf2* expression in whole tissue (n=7 pre-wean and n=5 adult) and (**f**) isolated SVCs (n=11 pre-wean – 12 adult). (**g**) Fluorescent *in situ* hybridization by RNAscope of *Igf2* (grey) in pre-wean and adult scWAT sections (n=3). Data shown by mean ± SEM.

### IGF2 does not influence the differentiation of primary preadipocytes *in vitro*

Both IGF1R and InsR signaling play important roles in adipogenesis. Thus, we investigated the role of *Igf2* in preadipocyte differentiation. As shown in **Fig. 1f**, adult preadipocytes express significantly lower levels of *Igf2* compared to pre-wean mice. Therefore, to mimic the pre-weaning conditions, primary adult preadipocytes were supplemented with 10 nM IGF2 throughout differentiation (d0-8) (**Fig. 2a**). *Igf2* mRNA levels remained the same in differentiated adipocytes compared to preadipocytes (**Fig. S2a**). Furthermore, supplementation of IGF2 did not alter the mRNA expression levels of adipogenic markers as well as protein levels of PPARγ (**Fig. 2b**, **Fig. S2b**). Additionally, lipid accumulation assessed by immunocytochemistry and Oil-Red-O (ORO) stainings did not show differences in lipid accumulation (**Fig. 2c and Fig. S2c**). We next tested the effects of inhibition of IGF2 on differentiation by treating primary pre-wean preadipocytes with 1 µg/mL IGF2 neutralizing antibody during the whole course of differentiation (**Fig. 2d**). The specificity of the antibody was further confirmed by treating the primary preadipocytes with either neutralizing antibody or IgG control for 24h in serum free media. The phosphorylation of AKT was reduced with the neutralizing antibody compared to the control implying that IGF2 signaling was blocked successfully (**Fig. S2d**). We observed a slight increase in the expression of adipogenic markers (*Ppar*γ, *AdipQ*, *Fabp4*) in cells differentiated with IGF2 neutralizing antibody compared to IgG controls. However, this did not reach statistical significance (**Fig. 2e**). The same trend was observed by morphological assessment using bright field imaging (**Fig. 2f**). In contrast to adult preadipocytes, *Igf2* expression was significantly reduced in differentiated adipocytes of pre-wean mice compared to preadipocytes (**Fig. S2e**) Lastly, we tested the role of IGF2 in adipogenesis by differentiating primary preadipocytes from adult scWAT through substitution of 100 nM insulin in the differentiation medium with either 10 nM IGF2 or 10 nM insulin. Cells treated with IGF2 differentiated significantly less compared to insulin treated cells as shown by *Ppar*γ and *Fabp4* expression (**Fig. 2g**). There was no significant difference in *Igf2* and *AdipQ* mRNA levels between conditions (**Fig. S2e**). These findings indicated that insulin is essential for preadipocyte differentiation, whereas IGF2 does not induce or enhance differentiation.

**Figure 2:**
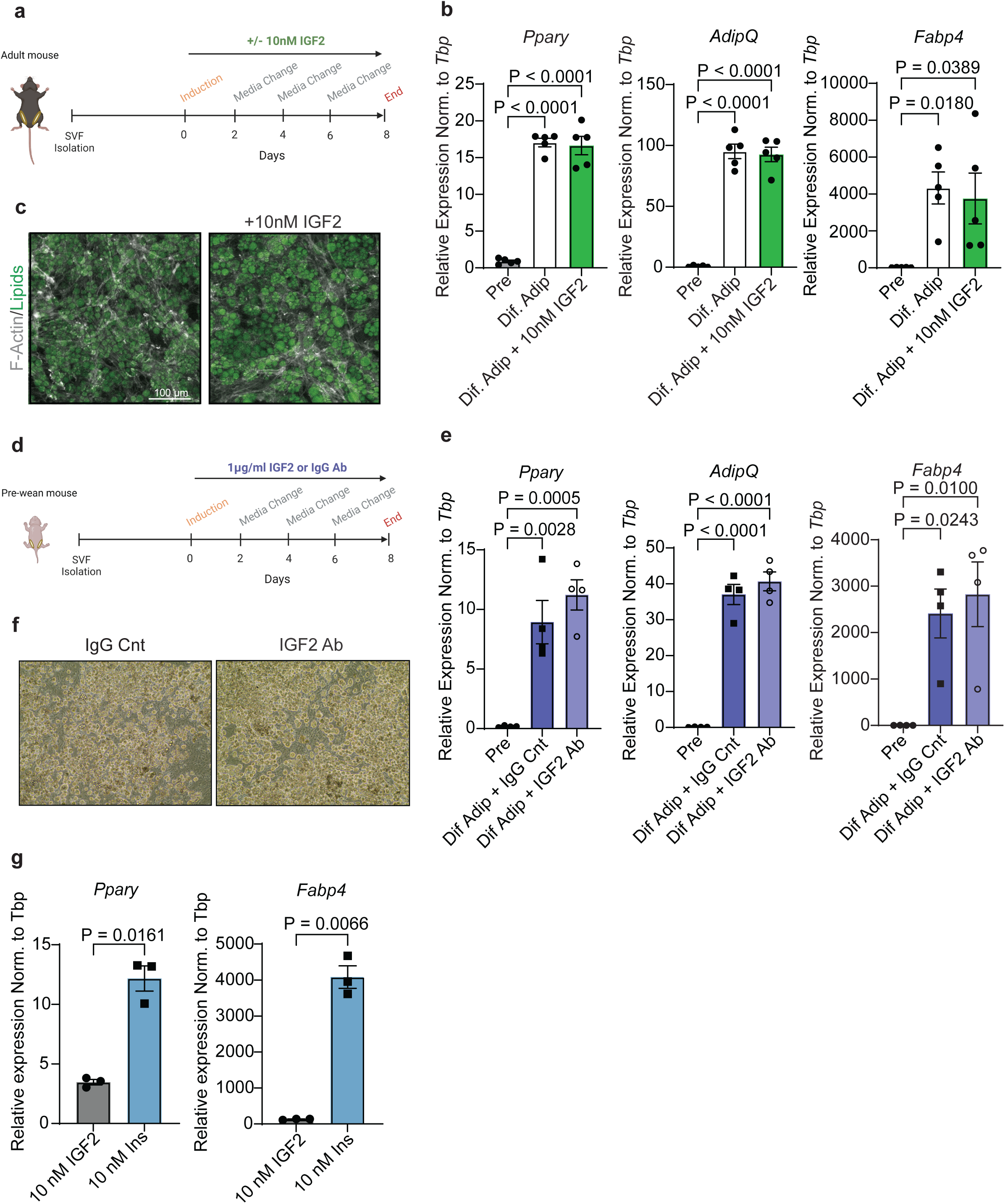
IGF2 supplementation does not affect differentiation of scWAT preadipocytes *in vitro*. (**a**) Schematic illustration of 10 nM IGF2 supplementation of cultured primary adult scWAT preadipocytes during the whole course differentiation. (**b**) mRNA levels of adipogenic markers (n=5). (**c**) Immunocytochemistry of lipids (green) and F-actin (grey) (n=3). (**d**) Schematic illustration of 1 µg/mL IgG or IGF2 neutralizing antibody (Ab) treatment of cultured primary pre-wean scWAT preadipocytes during the whole course differentiation. (**e**) mRNA levels of adipogenic markers (n=4). (**f**) Bright field images of differentiated adipocytes treated with either IgG or IGF2 neutralizing antibody (n=4). (**g**) 100 nM insulin was substituted with either 10nM IGF2 or insulin during differentiation. mRNA levels of adipogenic markers of differentiated primary adult preadipocytes with either IGF2 or insulin (n=3). Pre: preadipocytes, Dif. Adip: Differentiated Adipocytes. Data shown by mean ± SEM.

### IGF2 enhances proliferation of DPP4^+^ preadipocytes

IGF2 is a fetal growth hormone and highly abundant in fetal serum. To mitigate the potentially high endogenous level of IGF2 in FBS that may interfere with the effects of exogenously supplemented IGF2, we investigated alternative serum sources. Previous studies have shown that circulating IGF2 levels vary with age in mice. Consequently, our findings revealed that while serum insulin levels remained constant, IGF2 levels were significantly higher in pre-wean mice (239.4 ± 28.2 ng/mL) compared to adult mice (9.0 ± 0.6 ng/mL) (**Fig. S3a,b**). Additionally, pre-wean preadipocytes cultured overnight in serum-free media secreted slightly more IGF2 than adult preadipocytes, although this difference was not statistically significant (**Fig. S3c**). Based on these results, we proceeded to differentiate primary preadipocytes in media containing 1% adult mouse serum (MS). Interestingly, *Igf2* expression levels were significantly downregulated in adult preadipocytes differentiated in mouse serum (**Fig. S3d**). However, the differentiation capacity, as measured by mRNA levels of adipogenic markers, lipid content, immunocytochemistry of lipids, and protein levels of PPARγ, was unchanged in cells supplemented with 10 nM IGF2 compared to controls (**Fig. S3e-h**). A similar outcome was observed when pre-wean preadipocytes were treated with IGF2 neutralizing antibody compared to IgG controls and differentiated in mouse serum (**Fig. S3i,j**). Thus, despite the varying culture conditions, IGF2 does not directly influence adipogenesis.

Since IGF2 is a growth factor, we next tested whether it induces proliferation rather than differentiation. Primary adult preadipocytes were supplemented with 10 nM IGF2 for 7 days in 1% adult mouse serum. MTT assays revealed significantly higher proliferation rates of cells supplemented with 10 nM IGF2 compared to controls (**Fig. 3a**).

**Figure 3:**
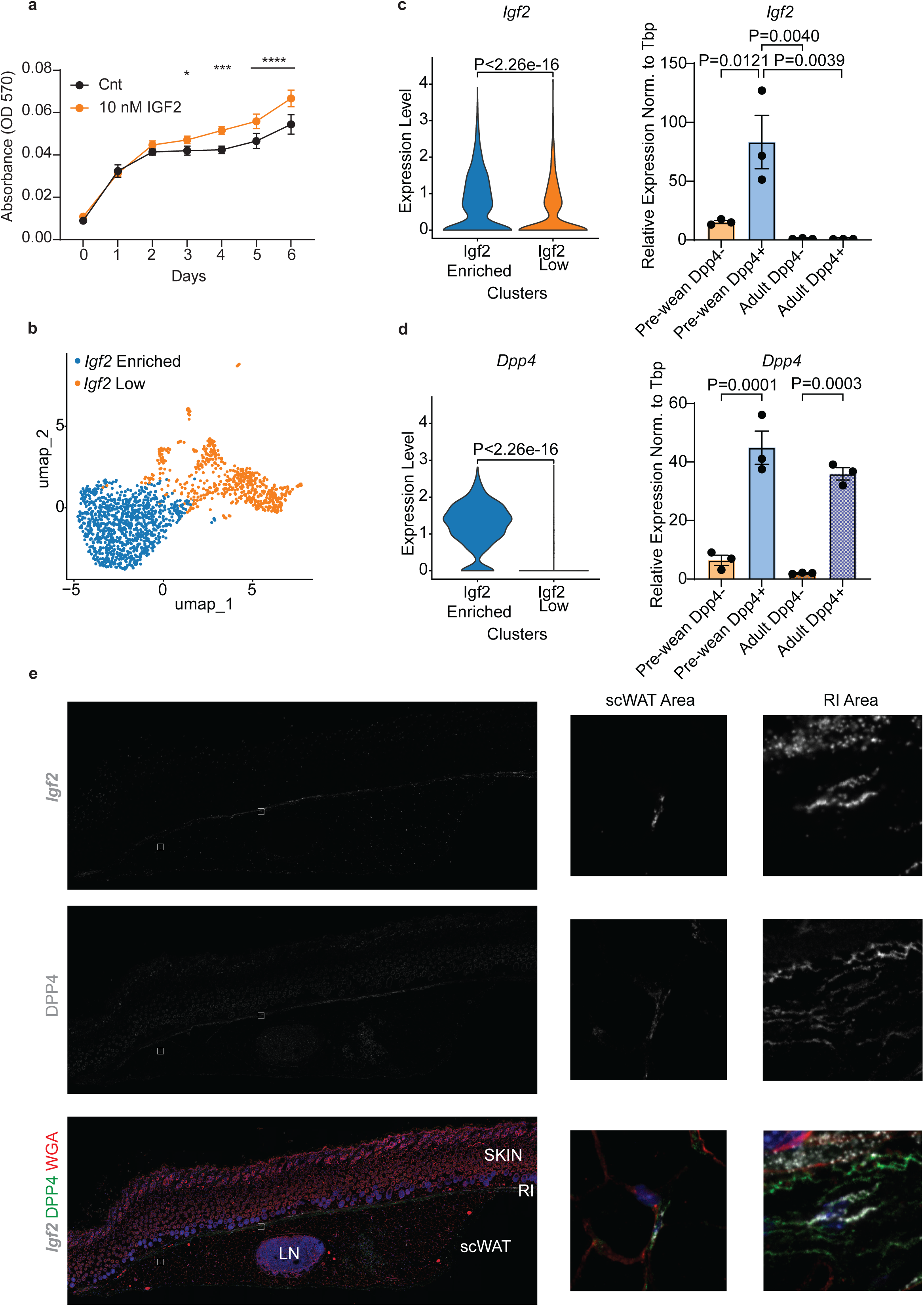
DPP4+ preadipocytes express significantly high levels of *Igf2*. (**a**) MTT assay of primary adult scWAT preadipocytes cultured with 1% adult mice mouse serum and supplemented with or without 10 nM IGF2 for 7 days (n=5). (**b**) Re-clustering of pre-wean preadipocytes only shown as UMAP plot. (**c**) Violin plot of *Igf2* expression between two clusters identified in (b) (left) and mRNA expression levels in DPP4 sorted cells from isolated pre-wean and adult SVCs (right) (n=3). (**d**) Violin plot of *Dpp4* expression between two clusters identified in (b) (left) and mRNA expression levels in DPP4 sorted cells from isolated pre-wean and adult SVCs (right) (n=3). (**e**) Fluorescent *in situ* hybridization by RNAscope of *Igf2* (grey) and immunohistochemistry of DPP4 (green), WGA (red) and DAPI (blue) was performed on pre-wean scWAT isolated with skin (n=4). The tile scan images were taken with 40x objective. * P=0.0222, ** P=0.0002, ****P<0.0001. Data shown by mean ± SEM.

To obtain more detailed insights into the pre-wean *Pdgfra^+^*preadipocytes populations expressing *Igf2,* we re-clustered these cells by *Igf2* enriched and low expressing cells (**Fig. 3b,c**). Differential gene expression analysis comparing *Igf2* enriched *vs* low cell clusters revealed 387 upregulated and 466 downregulated genes. Gene set enrichment analysis for Gene Ontology (GO) terms revealed that *Igf2* enriched cells express high levels of genes associated with ‘positive regulation of cell population proliferation’ and ‘extracellular matrix organization’, suggesting a potential role of *Igf2* in preadipocyte niche formation or maintenance (**Fig. S4a**). Furthermore, scRNAseq analysis revealed *Dpp4* as one of the most highly expressed genes in the *Igf2* enriched cell population. This finding was further validated by MACS sorting of DPP4^+^ preadipocytes, which showed significantly higher levels of *Igf2* expression compared to DPP4^-^ cells in pre-wean mice (**Fig. 3c-d**). In line with our previous results, *Igf2* expression was absent in adult DPP4^+^ cells (**Fig. 3d**). FISH and immunohistochemistry analysis showed that DPP4^+^*Igf2^+^*cells reside at the reticular interstitium (RI) region of pre-wean scWAT (**Fig. 3e**), which is critical for tissue expansion and regeneration (Merrick et al. 2019).

### Loss of *Igf2* promotes differentiation of DPP4^+^ SVCs

Given our data indicating that IGF2 increases cell proliferation and prior research characterizing DPP4 preadipocytes as highly proliferative (Merrick et al. 2019), we explored whether *Igf2* helps maintain proliferation of DPP4^+^ cells early in life. Analysis of the scRNAseq data from pre-wean DPP4^+^ cells revealed higher expression of the proliferation marker *Ki67*, compared to adult DPP4^+^ cells (**Fig. 4a**). qPCR analysis did not show a general difference in *Ki67* expression between pre-wean and adult DPP4^+^ SVCs, while we observed a significant difference between pre-wean DPP4^+^ and DPP4^-^ cells (**Fig. 4b**). Furthermore, we found that pre-wean DPP4^+^ cells proliferated at a higher rate than adult cells (**Fig. 4c**). Moreover, *Dpp4*^+^*Igf2*^+^ cells express higher *Ki67* levels compared to *Dpp4*^+^*Igf2*^-^ (**Fig. 4d**). These findings suggest that *Igf2* expression promotes proliferation of DPP4^+^ cells and prompted us to hypothesize that its loss-of-function would commit DPP4^+^ cells to differentiation. To test this, *Igf2* was knocked down (siIgf2) in primary DPP4^+^ cells from pre-wean scWAT using small interference RNA (siRNA). Evaluation of adipogenic markers by qPCR and bright field images revealed that siIgf2 preadipocytes differentiated significantly more compared to control cells (siNeg) (**Fig. 4e,f**). These findings strongly support our finding that *Igf2* is essential for DPP4^+^ cells to maintain their proliferative capacity and precursor nature early in life.

**Figure 4:**
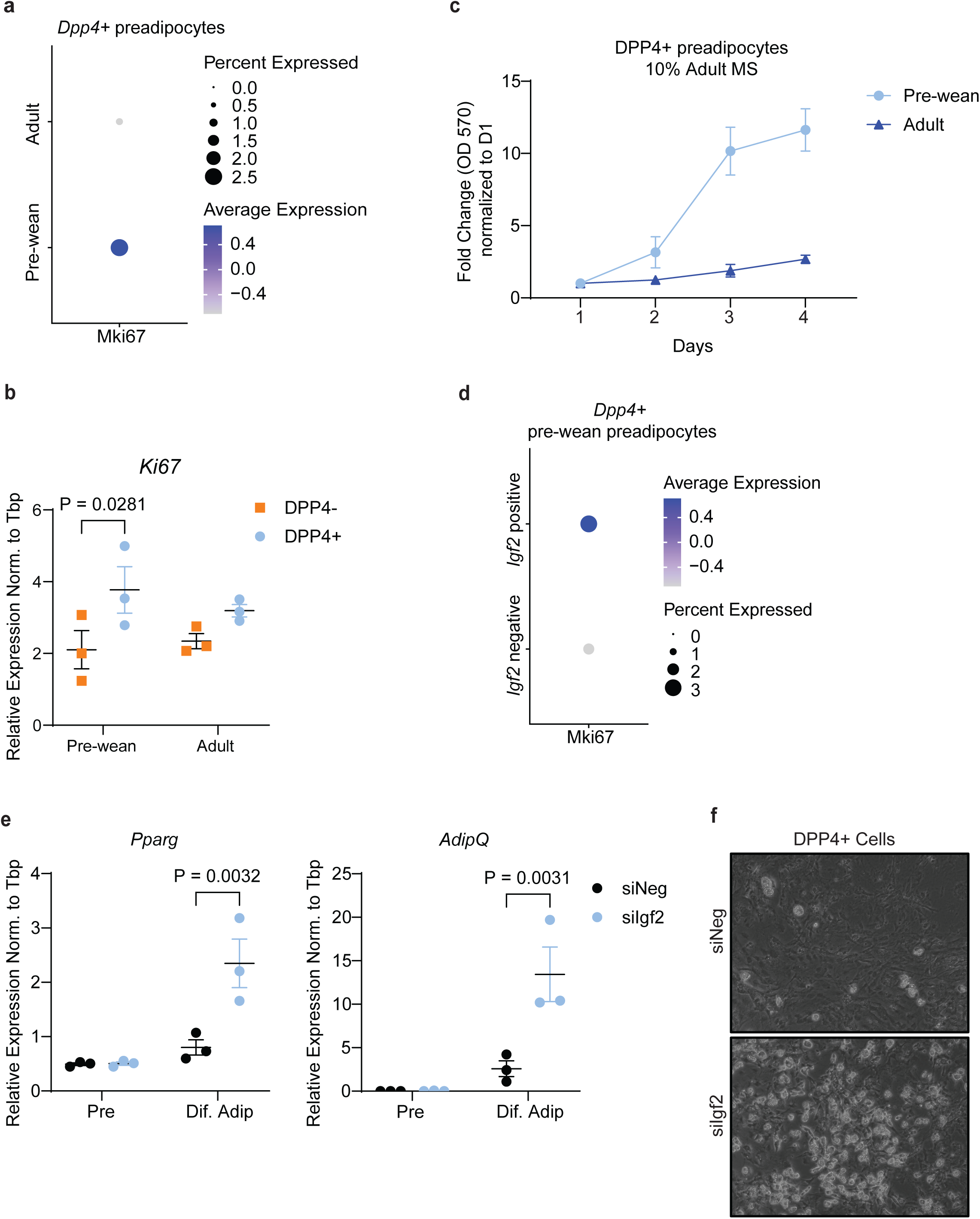
Loss of *Igf2* in DPP4+ cells induce preadipocyte differentiation. (**a**) Dot plot of *Ki67* expression on pre-wean and adult *Dpp4*+ preadipocytes based. (**b**) mRNA expression of *Ki67* in MACS sorted DPP4+ and DPP4-pre-wean and adult SVCs (n=3). (**c**) MTT assay of DPP4+ primary preadipocytes isolated from pre-wean and adult mice scWAT (n=2). (**d**) Dot plot of Ki67 in *Dpp4*+ pre-wean preadipocytes comparing *Igf2* positive to negative cells. *Igf2* was knockdown (siIgf2) in pre-wean DPP4+ primary preadipocytes. (**e**) mRNA expression of adipogenic markers (n=3). (**f**) Bright field images of *Igf2* knockdown and control differentiated pre-wean preadipocytes (n=3). Data shown by mean ± SEM.

Adipose tissue development and expansion occur via hyperplasia and hypertrophy (Wang et al. 2013). Subcutaneous adipocyte hyperplasia is associated with healthier metabolism (Vishvanath and Gupta 2019). This process is tightly regulated by preadipocyte proliferation and *de novo* adipogenesis. Since preadipocyte populations tightly regulate this process, it is critical to understand different subtypes present in adipose tissue and the microenvironment they create to regulate tissue expansion (Altun et al. 2022).

To this end, we previously described scRNAseq analysis on scWAT SVCs comparing pre-wean and adult mice, aiming to identify compositional differences during developmental states that positively regulate adipose tissue homeostasis. In this study, we re-analyzed this dataset focusing on *Pdgfra^+^* preadipocytes, as the most distinct separation by age was observed in this cell population (Suwandhi et al. 2021). As a result, we identified *Igf2* as the most abundantly expressed gene in pre-wean scWAT preadipocytes. This was further confirmed at mRNA levels in whole tissue and isolated SVCs, and by FISH. The significant decrease of *Igf2* in adult mice suggests a critical developmental function of the gene in adipose tissue development. Additionally, even though IGF2 is an important fetal growth promoter, the postnatal role of the protein is not well understood. Consistent with previous data, we find that 10 nM IGF2 activates PI3K/AKT and MAPK/ERK pathways through IGF1R and InsR in primary preadipocytes (Frasca et al. 1999; Blyth et al. 2022). It was previously shown that loss of both receptors in preadipocytes severely affects differentiation (Boucher et al. 2010).

First the role of IGF2 in adipocyte differentiation was investigated by supplementing adult preadipocytes with IGF2 to mimic the high *Igf2* expression in pre-wean mice, whereas pre-wean preadipocytes were treated with neutralizing antibody to block receptor-ligand interaction. Even though blockage of IGF2 and receptor interaction showed a slight increase in mRNA levels of adipogenic markers, there were no significant differences between control and treatment conditions in both approaches.

A study on normal weight children showed that IGF2 supplementation enhances scWAT preadipocyte differentiation while it inhibits visceral preadipocyte differentiation in a dose-dependent manner (7.5 and 62.5 ng/mL) (Alfares et al. 2018). Additionally, rat adipose stem cells showed that supplementation of 100 ng/mL IGF2 only under 1% FBS promotes adipocytes differentiation and self-renewal capacity through IGF1R and InsR but had no effect using 10% FBS (Wang et al. 2019). Differences between previous studies and our results could be due to species differences or the fact that white adipose tissue SVCs are heterogenous and cellular composition of isolated preadipocytes can vary between studies based on age and isolation site. This has been discussed previously (Altun et al. 2022; Palani et al. 2023). Moreover, since IGF2 is a fetal growth factor, exogenous protein levels present in fetal serum could impact on study results. To overcome these limitations and uncertainties, we substituted FBS with adult mouse serum, that contains low levels of IGF2. We did not observe differences in the differentiation of preadipocytes between treatment and control groups both in pre-wean and adult cells. Along with these findings, substitution of insulin in the differentiation media with IGF2 significantly reduced mature adipocyte gene expression and differentiation of preadipocytes. Thus, our data strongly support the conclusion that IGF2 does not enhance adipogenesis of primary murine preadipocytes.

Several studies have highlighted the role of IGF2 promoting proliferation and expansion of various cell types (Wang et al. 2018; Wang et al. 2019; Sandovici et al. 2022). In line with this, our results demonstrated that supplementation of adult preadipocytes that express low levels of *Igf2* with 10 nM IGF2 enhanced their proliferation. scRNAseq analysis revealed that *Igf2^+^*cells express significantly high levels of *Dpp4* early in life. Consistent with the results above, these cells lost *Igf2* expression in adulthood. Merrick *et al*. identified DPP4^+^ cell population as highly proliferative and multipotent precursors, however, the mechanism behind this is not well understood (Merrick et al. 2019). MTT assays showed that isolated adult DPP4^+^ cells proliferate significantly less compared to pre-wean. We showed that one of the significant differences between these two cell populations is the expression of *Igf2*. However, we cannot exclude additional physiological changes during development. Thus, to better evaluate the function of *Igf2* specifically in DPP4^+^ cells, we performed a loss of function study with small interference RNA (siRNA) in sorted pre-wean DPP4^+^ preadipocytes. *Igf2* knockdown cells expressed significantly higher levels of adipogenic markers and showed more lipid accumulation compared to control knockdown. The same phenomenon has been reported with knockdown of *Dpp4*, which induced human preadipocyte differentiation (Hatzmann et al. 2022). These findings demonstrate that *Dpp4* and *Igf2* expression regulate proliferation and expansion of DPP4^+^ precursor population and loss of expression of either one of them commits cells to differentiation.

In conclusion, we show for the first time a key function of *Igf2* expression in a specific adipose precursor cell population. Our findings suggest that *Igf2* expression maintains the proliferative capacity of DPP4^+^ cells early in life to serve as a reservoir for adipose progenitor pool. Thus, our study allows a better understanding of developmental mechanisms in adipose progenitor populations, found in a distinct location, to foster new therapeutic approaches for promoting metabolically healthy adipose tissue expansion.

## Methods

### Mice

Pre-wean (15 ± 2 days old) and adult (58 ± 2 days old) C57BL/6 wild type mice were housed at constant ambient temperature of 22 ± 2°C, 45-65% humidity and a 12h light-dark cycle with *ad libitum* access to standard chow diet (Altromin 1314, Lage Germany) and water. During the sacrifice of animals, blood was withdrawn and kept at room temperate (RT) for 15 mins prior to incubation on ice. Mouse serum (MS) was obtained by centrifugation at 10,000 x g for 5 min at RT. Serum from 27 mice was pooled, filtered through a 0.45 µm filter and stored at - 80°C for cell culture experiments. All animal procedure were performed under the guidance and approval of German animal welfare law and district government of Upper Bavaria (Bavaria, Germany).

### Primary cell culture

Primary cells were isolated from subcutaneous adipose tissue (scWAT) of pre-wean and adult C57Bl6 wild type male mice. Freshly isolated tissues were chopped into small pieces and digested in digestion solution (Dulbecco’s modified Eagle’s medium high glucose + GlutaMAX (DMEM) (Gibco) containing 1% BSA and 1 mg/mL collagenase Type I (Life Technologies)) at 37 °C and 1,000 x rpm shaking for 30-40 min. Collagenase Type I was substituted by collagenase Type IV (Life Technologies) to isolate DPP4+ preadipocytes. The suspension was then filtered through a 100 µm strainer. Cells were centrifuged at 500 x g for 5 min at RT. Pelleted SVCs were washed one time with PBS and either stored at -80°C for total RNA isolation or cultured in normal growth media (DMEM, 10% fetal bovine serum (FBS), 1% penicillin and streptomycin (PenStrep) and 0.1 mg/mL Normocin (Invitrogen) at 37°C and 5% CO_2_.

#### Preadipocyte Differentiation

Confluent preadipocytes were induced (day 0, d0) in DMEM with 10% FBS or Opti-MEM with 1% MS containing 0.5 mM 3-isobuthyl-1-methylxanthin (IBMX, Sigma Aldrich), 5 μM dexamethasone (D4902, Sigma Aldrich), 100 nM insulin (Sigma Aldrich), 1% PenStrep and 0.1 mg/mL Normocin. After 2 days, the induction medium was replaced by differentiation medium with only 100 nM insulin in respective culture media conditions. The media was changed every two days until day 8 (d8) of differentiation. In the case of supplementation or neutralization experiments, cells were cultured with 10 nM IGF2 (Sigma-Aldrich, I8904) or 1 μg/mL IGF2 neutralizing antibody (IGF2 Ab, Thermo Scientific, PA5-47946), respectively, from d0 to d8 of differentiation. Goat IgG (1 μg/mL, Invitrogen, 31245) was used as a control for the blocking experiments and no treatment was used as a control for supplementation experiment. Adult scWAT preadipocytes were differentiated by substituting 100 nM insulin with 10 nM IGF2 or 10 nM insulin from d0 to d8. IBMX and dexamethasone were still present between d0 and d2 of differentiation. At the end of the experiment cells were either fixed or frozen for further analysis.

#### Igf2 knock down

Cultured DPP4+ preadipocytes were transfected following a reverse transfection method (Isidor et al. 2016). Briefly, preadipocytes were trypsinized and collected in culture media. A transfection mix was prepared mixing OptiMEM (Gibco), Lipofectamine RNAiMAX (Invitrogen) and 50 nM siRNA and incubating 20 min at RT. Cells were then seeded with the transfection mix and media was changed after 24h. Cell were grown to confluence and induced with 0.5 mM IBMX, 5 μM dexamethasone, 100 nM insulin. Induction media was changed to differentiation medium containing 100 nM insulin. The media was changed every 2 days for 6-7 days. The *Igf2* siRNA (siIgf2) sequence was modified from Gui et al. (Gui et al. 2021) (mmIgf2 Frw: 5’GGAGCUUGUUGACACGCUUCAGUU3’ Rev: 5’CCUCGAACAACUGUGCGAAGUCAA3’) (IDT). A commercially available negative control siRNA (siNeg) was used (51-01-14-04, IDT).

### Single cell RNA (scRNA) sequencing and analysis

scRNA sequencing data were previously published (Suwandhi et al. 2021) and further analyzed using R. The Cell Ranger single cell software suite v2.1.1 from 10X Genomics was used to preprocess the data. This included demultiplexing raw BCL files to fastq files, mapping of reads to an indexed mm10 reference genome, which was built using the GRCm38 assembly with ensembl genome annotation release 94 and UMI counting. A gene expression count matrix along with feature barcode counts for each cell barcode was generated resulting in 20151 genes across 9319 cells. These matrices were then processed in R (v4.3.2) using the Seurat package (v5.1.0). The initial QC steps included filtering out cells which captured <500 expressed genes and potential doublets with a total number of detected genes >5000. The percentage of mitochondrial genes expressed were computed and cells with more that 10% mitochondrial genes were discarded. The remaining cells were normalized and scaled using default parameters of Seurat functions.

For cluster generation, we performed PCA and based on elbow plot top 20 PCs were considered to generate a UMAP. Cell type was determined by examining the expression of a set of marker genes that were reviewed from the literature and observed along the clusters. Based on interested cell-types and batch, further subset of clusters followed by coarser clustering were carried out. We considered preadipocytes clusters for the pre-wean mice to check expression of *Igf2* and *Dpp4* positive cells. For the volcano plot, differentially expressed genes were generated using FindMarkers function of Seurat by assigning the batch type comparisons as idents. Violin plots, dotplots and individual UMAP plots for the specific genes were performed by Seurat’s VlnPlot, Dotplot and FeaturePlot functions, respectively. We predicted classification of each cell in either G2M, S or G1 phase by assigning scores using the CellCycleScoring function. Gene Ontology (GO) analysis was performed in R using the package enrichR.

### MACS sorting of DPP4+ cells

Freshly isolated SVCs from scWAT of pre-wean and adult wild type male mice were washed with PBS and blocked in 1% BSA and 1mM EDTA in PBS for 10 mins at 4°C. Four to six mice were pooled for each biological replicate. Cells were incubated with a DPP4 antibody (1:10, R&D Systems, AF954) for 30 min, followed by addition of 15 µL protein G magnetic beads (Milteny Biotech) for 30 min. MS or LS columns (Milteny Biotech) were calibrated with blocking buffer and loaded with cells. The flow through fraction was collected for the assessment of DPP4^-^ cells. The column was then washed three times with blocking buffer, removed from the magnetic board and DPP4^+^ cells were eluted in 500 µL blocking buffer. Cells were either pelleted and kept in RLT-DTT buffer for mRNA assessment or cultured in normal growth media for future experiments.

### RNA isolation and RT-qPCR

Whole tissue samples were lysed for 2 min at 30 Hz/sec in 1 mL Qiazol (Qiagen) with metal beads in the TissueLyzer II (Qiagen). Samples were kept at RT for 5 min, 200 µL Chloroform was added and samples were incubated another 3 min at RT. Homogenates were centrifugated at 12,000 x g for 15 min at 4°C. Afterwards the clear aqueous phase was transferred to a fresh tube and mixed with the same volume of 70% ethanol and transferred to RNAeasy mini spin columns. For RNA isolation from preadipocytes and differentiated adipocytes, cell pellets were lysed in RLT buffer containing 25 µM dithiothreitol (DTT). Samples were then mixed with the same volume of 70% ethanol and transferred to RNAeasy mini spin columns. RNA isolation was performed following the manufacturers recommendations using a RNAeasy Mini Kit (Qiagen), except for DPP4 sorted cells, where an RNAeasy Micro Kit was used. RNA yield was determined using aNanoDrop 2000 UV-Vis Spectrophotometry (Thermo Scientific).

To determine gene expression, cDNA (300-500 ng total RNA) was synthesized using the High-Capacity cDNA Reverse Transcription Kit (Applied Biosystems). Relative mRNA expression was quantified by mixing cDNA, 300 nM of forward and reverse primers and iTaq Universal Green Supermix (Bio-Rad), running in a C1000 Touch Thermal Cycler (Bio-Rad). Samples were run in duplicates or triplicates for each gene and quantified using the Biorad CFX Manager 3.1 Program. All expression levels were normalized to TATA-binding protein (*Tbp*). Cq value was set to 50 when there was no gene amplification, but *Tbp* expression was detected. Primer sequences used for the analysis were as follows:

*Igf2* Fwd: 5’AGACATACTGTGCCACCCC3’ Rev: 5’ATTGGAAGAACTTGCCCACG3’,

*Fabp4* Fwd: 5’GATGCCTTTGTGGGAACCT3’ Rev: 5’CTGTCGTCTGCGGTGATTT3’,

*Ppar*γ Fwd: 5’CCCTGGCAAAGCATTTGTAT3’ Rev: 5’GAAACTGGCACCCTTGAAAA3’,

*AdipQ* Fwd: 5’GATGGCACTCCTGGAGAGAA3’ Rev: 5’TCTCCAGGCTCTCCTTTCCT3’,

*Tbp* Fwd: 5’ACCCTTCACCAATGACTCCTATG3’ Rev: 5’TGACTGCAGCAAATCGCTTGG3’,

*Dpp4* Fwd: 5’TCAGCTCATCCTCTAGTGCG3’ Rev: 5’AGCCCACACCACATCACATA3’,

*Ki67* Fwd: 5’CCTTTGCTGTCCCCGAAGA3’ Rev: 5’GGCTTCTCATCTGTTGCTTCCT3’.

### Protein isolation and western blot

Differentiated preadipocytes or cultured primary pre-wean preadipocytes that were serum starved for 24 h in the presence of 1 μg/mL IgG or IGF2 neutralizing antibody were washed with ice cold PBS and lysed using RIPA buffer (50 mM Tris pH 7.4, 150 mM NaCl, 1mM EDTA, 1% Triton X-100) containing 0.1% sodium dodecyl sulfate (SDS), 0.01% protease inhibitor, 0.01% phosphatase inhibitor cocktails II and III (all Sigma). Lysates were incubated on ice for 10 min and centrifuged for 10 min at 14,000 x g at 4°C. Protein concentration was determined by BCA Protein Assay Kit (Thermo Scientific). Samples were mixed with NuPAGE Sample Buffer (Life Technologies) containing 2.5% beta-mercaptoethanol (Carl Roth) and heated at 95°C for 5 min. Samples were then loaded on an SDS-PAGE gel with a Fischer BioReagents EZ-Run Prestained Rec Protein Ladder (Thermo Scientific) and transferred to PVDF membranes (0.45 µm, Merck Millipore). Unspecific binding sites were blocked using 5% bovine serum albumin (BSA) or non-fat dried milk in TBS-T (1 X TBS containing 0.1% Tween-20) for 1h at RT, followed by incubation with primary antibodies overnight at 4°C: AKT (1:1000, Cell Signaling, 4685), phosphor-AKT Ser 473 (1:1000, Cell Signaling, 9271), ERK (1:1000, Cell Signaling, 4695), phosphor-ERK (1:1000, Cell Signaling, 4377), insulin receptor (InsR) (1:1000, Cell Signaling, 3020), IGF1 receptor (IGF1R) (1:1000, Cell Signaling, 9750), PPARγ (1:1000, Cell Signaling, 2443), B-Actin HRP (only 30 min at RT, 1:5000, Santa Cruz, Sc-47778). Next day, membrane was washed with TBS-T and incubated with secondary antibodies 1 h at RT: anti-rabbit HRP (1:5000, Cell Signaling, 7074) and anti-mouse HRP (1:5000, Santa Cruz, Sc-2005). After washing, the membrane was incubated with either ECL (Merck Millipore) or SuperSignal ECL (Thermo Scientific) for 1 min and imaged by the ChemiDoc (Bio-Rad).

#### Immunoprecipitation (IP)

Cultured primary scWAT preadipocytes were serum starved for 3 h and stimulated with 10 nM IGF2 for 10 min and lysed in IP lysis buffer (RIPA with 2 mM EDTA, 0.01% protease inhibitor, 0.01% phosphatase inhibitor cocktails II and III). Protein isolation was the same as mentioned above. Two hundred microgram protein lysates were incubated with either anti-InsR or anti-IGF1R antibodies (both 1:100) overnight at 4°C with gentle rotation. Next day, lysate-antibody mixes were incubated with 15 μL A/G agarose beads for 1-2h at 4°C and washed 3 times with IP lysis buffer. The immune complex was eluted by incubation the lysates with 2 X NuPAGE Sample Buffer (Life Technologies) containing 5% beta-mercaptoethanol and boiling at 95°C for 5 min. Samples were run on a NuPAGE 4-12% Bis-Tris Gel (Invitrogen) in 1 X MOPS Buffer at 200V and transferred to 0.45 μm PVDF membranes (Merck Millipore). The membrane was blocked with 5% BSA and incubated with primary antibodies overnight at 4°C: phospho-Tyrosine (1:1000, Cell Signaling, 8954), InsR (1:1000, Cell Signaling, 3020), IGF1R (1:1000, Cell Signaling, 9750), B-Actin HRP (only 30 min at RT, 1:5000, Santa Cruz, Sc-47778). The next day, the membranes were washed with TBS-T and incubated with secondary antibodies for 1 h at RT: anti-rabbit HRP (1:5000, Cell Signaling, 7074) and anti-mouse HRP (1:5000, Santa Cruz, Sc-2005). After the washing steps, the membrane was incubated with SuperSignal ECL (Thermo Scientific) for 1 min and imaged using the ChemiDoc (Bio-Rad).

### MTT Assay

Isolated scWAT preadipocytes were grown to 80% confluence and seeded on 96-well plates with 1 x 10^4^ cells per well. A plate was prepared for each day of the measurement. Cells were incubated with 0.5 mg/mL MTT final concentration in the well for 2 h at 37°C, except on the seeding day cells were incubated 3 hours post seeding with MTT. Death control wells were treated with 0.03% Triton X-100 for 1 min and then MTT was added to the wells. Formazan was eluted with solubilization solution (10% Triton X-100, 0.03% HCl in 100% isopropanol) shaking at 700 rpm for 10 min at RT. The supernatant was then transferred into a fresh 96-well plate (Greiner, F-bottom) and its absorbance was measured at 570 nm using either a Varioskan Lux Multimode plate reader (Thermo Scientific) or a PHERAstar FSX (BMG Labtech). Background absorbance was measured at 640 nm. The preadipocytes of adult mice were supplemented with 10 nM IGF2 starting from the first day of seeding until day 7 cultured in 1% Adult MS (d0-7).

### Fluorescent *in situ* hybridization by RNAscope

RNAscope staining (Bio-techne, 323133) of mm-Igf2 (Atto 570, Biotechne, 437671) was performed on 20 µm cryosections of pre-wean and adult scWAT according to the manufacturer’s instructions. Two µm paraffin sections of pre-wean scWAT with skin were stained for mm-Igf2 and TSA Vivid Fluorophore 570 (Bio-techne, PN 323272) following the manufacturers manual of RNAscope (Bio-techne, 323100), except the antigen retrieval was for 30 min. RNAscope 3-plex Positive (Bio-techne, 320881) and Negative (Bio-techne, 320871) probes were used as control stainings. Sections were subsequently blocked with 10% goat serum for 1h at RT and stained for DPP4 (1:1000, Abcam, ab187048) in 5% goat serum overnight at 4°C. Slides were washed three times with PBS for 5 min in RT and stained with secondary antibodies Donkey anti Rabbit Alexa 488 (1:400, Invitrogen, A21206) and WGA conjugated Alexa 647 (1:1000, Invitrogen, W32466) for 1h at RT. After washing the slides three times with PBS for 5 min, sections were incubated with DAPI for 30 sec and mounted with DAKO mounting media. Fluorescent signal was imaged using SP8 Confocal microscope (Leica).

### ELISA

Serum levels of insulin and IGF2 were measured using ultra-sensitive mouse insulin ELISA kits (90080, Crystal Chem) and m/r/p/calIGF-II Qkit (MG200, R&D Systems), respectively, following the manufacturer’s instructions.

### Immunocytochemistry

*In vitro* differentiated primary adipocytes were fixed with 10% formalin for 10 min at RT, permeabilized with 0.1% Triton X-100 on ice and blocked with 3% BSA containing 0.03% Triton X-100 for 1h at RT. Slides were stained for Alexa 647 conjugated Phalloidin (1:40, Life Technologies, A22287) and Lipitox Green (1:200, Life Technologies, H34475) in blocking buffer for 1h at RT and washed with three times PBS. Sections were mounted using Dako fluorescence mounting medium, and images were acquired using a SP8 Confocal microscope (Leica).

### Oil Red O (ORO) staining

*In vitro* differentiated primary preadipocytes were fixed with 10% formalin at RT. Cells were dehydrated with 60% isopropanol for 5 min and dried. Cells were incubated with ORO working solution (60% stock diluted in ddH_2_O) for 10 min at RT and washed with ddH_2_O. Cell number was measured after staining cells with DAPI (1:5000) for 5 min and measuring fluorescence intensity at 460 nm using PHERAstar FSX (BMG Labtech). ORO was diluted in 100% isopropanol and absorbance was measured at 500 nm using PHERAstar FSX.

### Statistical Analysis

Statistical significance for multiple comparisons was determined by ordinary One-way ANOVA or Two-way ANOVA, with Tukey’s multiple comparisons test, or unpaired Two-Tailed t-test using GraphPad prism 10.0.2 program. All the statistical tests for the scRNAseq experiments were performed in R, the barplots were generated using the ggplot2 package. All the respective p-values in violin plots are calculated using the Wilcoxon test and stat_compare_means function from the ggpubr package for respective condition pairs. Data are shown as ± standard error mean (SEM). Exact P-values were indicated in the figures and P<0.05 was considered as statistically significant.

## Acknowledgments

We would like to thank all members of the Ussar lab for fruitful discussions. S.W. and F.P. were supported through the Initiative and Network Fund of the Helmholtz Association, and the Munich Center for Systems Neurology (SyNergy EXC 2145; Project ID 390857198).

## Author Contributions

I.A. designed and conducted experiments and wrote the manuscript. S.W., V.M., D.L and F.P. conducted the scRNAseq data analysis, data interpretation and contributed to writing the manuscript. X.Y., I.S., R.K., L.K. and KHDV conducted experiments and contributed to writing the manuscript. S.U. designed the study, coordinated the experiments and wrote the manuscript.

---

**Supplementary Figure 1:**
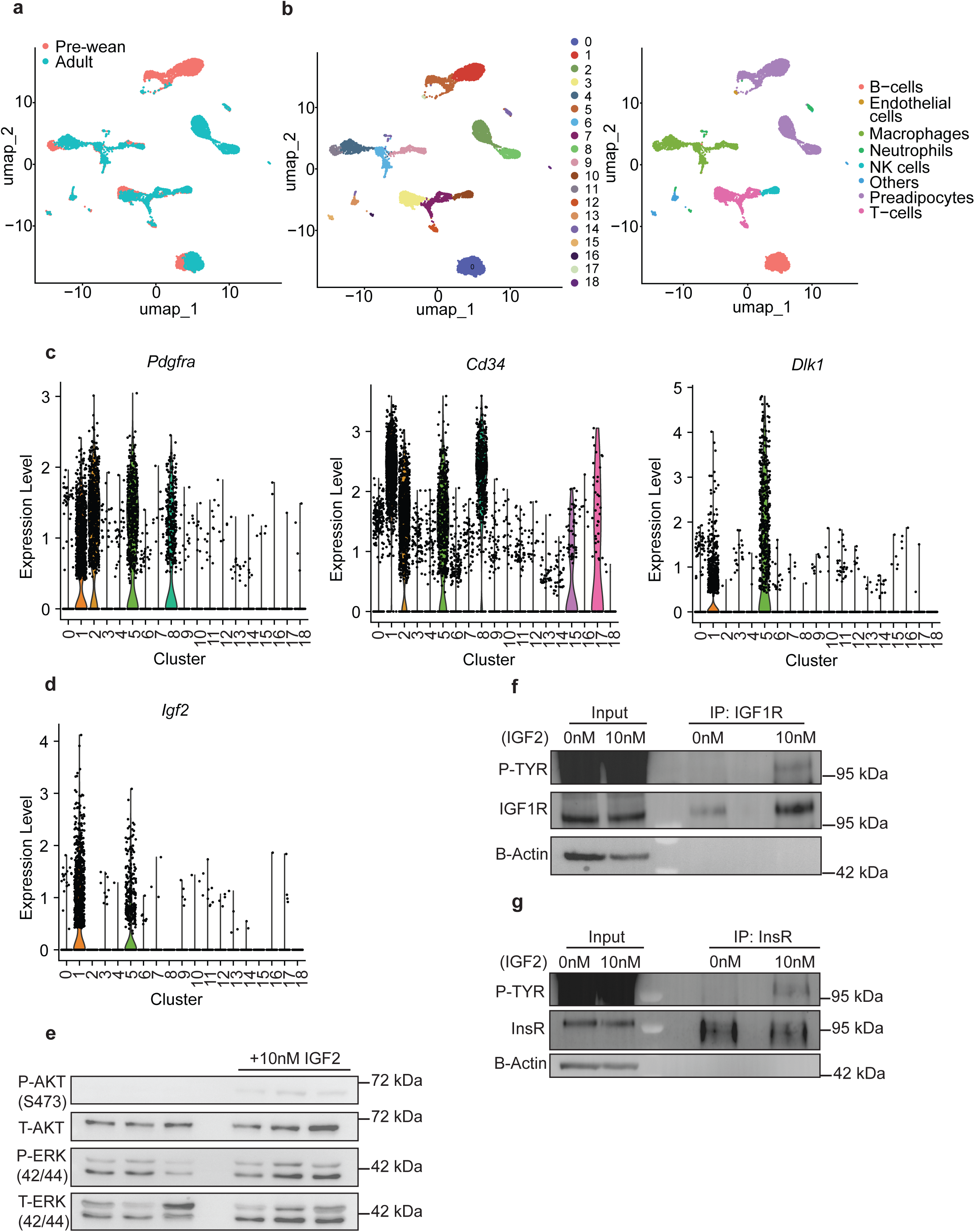
IGF2 activates AKT and ERK through IGF1R and InsR signaling pathways. (**a**) Projection of two age groups (pre-wean and adult) on isolated scWAT SVCs shown as UMAP plot. (**b**) Cell Clustering of (a) based on cell types. Violin plot of (**c**) preadipocyte markers and (**d**) *Igf2*. Adult primary scWAT preadipocytes were serum starved 3h and stimulated with 10 nM IGF2 for 10 min. (**e**) Western blot of phosphorylated (P)/ total (T) AKT and ERK (n=3). Immunoprecipitation (IP) of (**f**) IGF1R (n=2) and (**g**) InsR (n=2). Immunoblotting of P-Tyrosine (TYR) was used to assess the phosphorylation of the receptors after IGF2 stimulation.

**Supplementary Figure 2:**
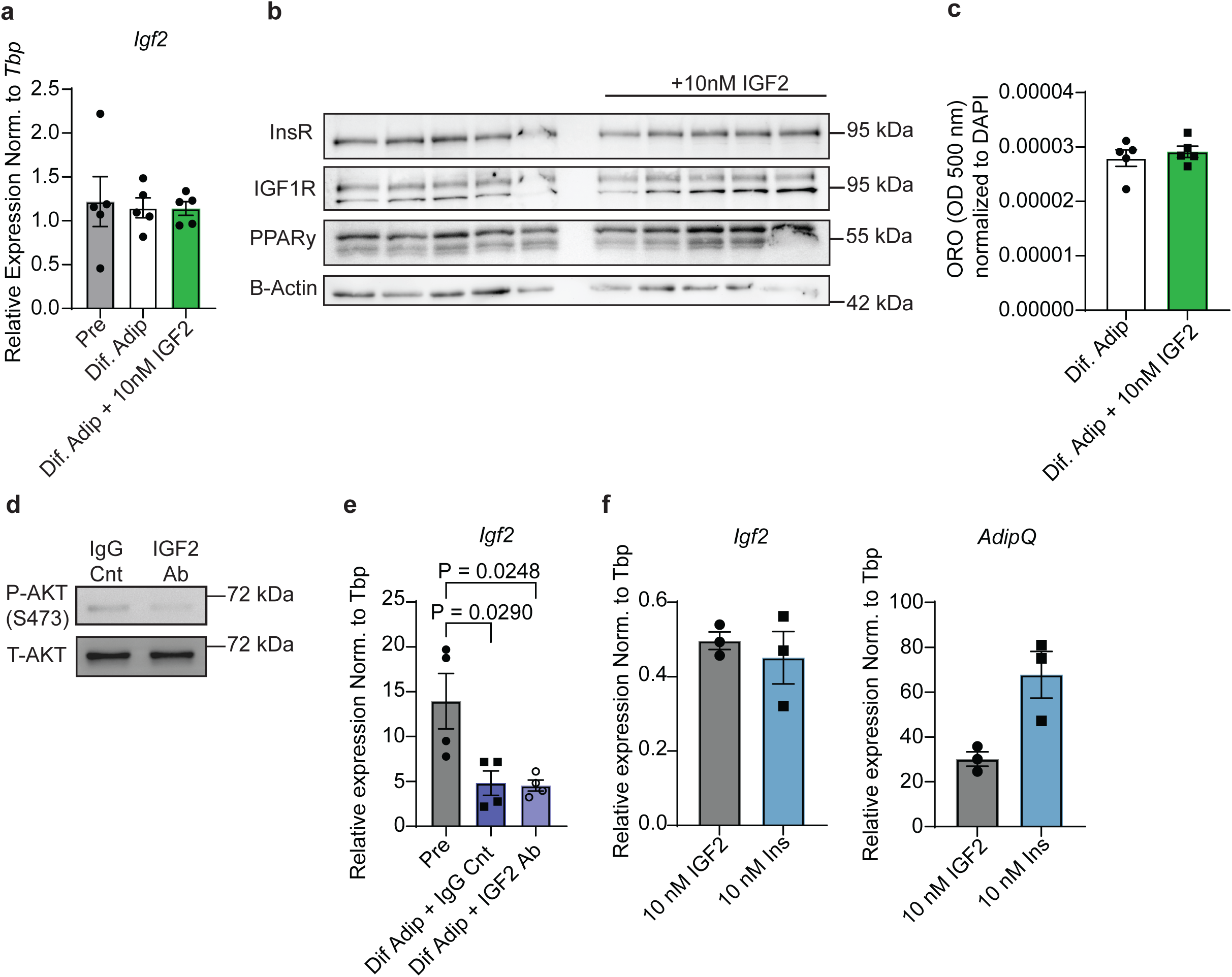
Cultured adult scWAT preadipocytes (Pre) were differentiated with (Dif. Adip + 10nM IGF2) or without (Dif. Adip) 10 nM IGF2. (**a**) mRNA expression of *Igf2* (n=5). (**b**) Western blot of InsR, IGF1R and PPARγ. B-Actin was used as loading control (n=5). (**c**) Lipid quantification by Oil red O (ORO) absorbance at 500 nm normalized to DAPI of only differentiated primary adult scWAT preadipocytes (n=5). (**d**) Western blot of phosphorylated (P) and total (T) AKT of protein isolated from cultured pre-wean scWAT preadipocytes that were serum starved for 12 h with 1µg/ml IgG or IGF2 neutralizing antibody (IGF2 Ab) (n=1). (**e**) mRNA levels of *Igf2* from differentiated pre-wean scWAT primary preadipocytes that were treated with 1 µg/ml IgG (Dif. Adip + IgG Cnt) or IGF2 neutralizing antibody (Dif. Adip IGF2 Ab) (n=4). (**f**) 100 nM insulin was substituted with either 10 nM IGF2 or insulin during differentiation. mRNA levels of *Igf2* and *AdipQ* (n=3). Data shown by mean ± SEM.

**Supplementary Figure 3:**
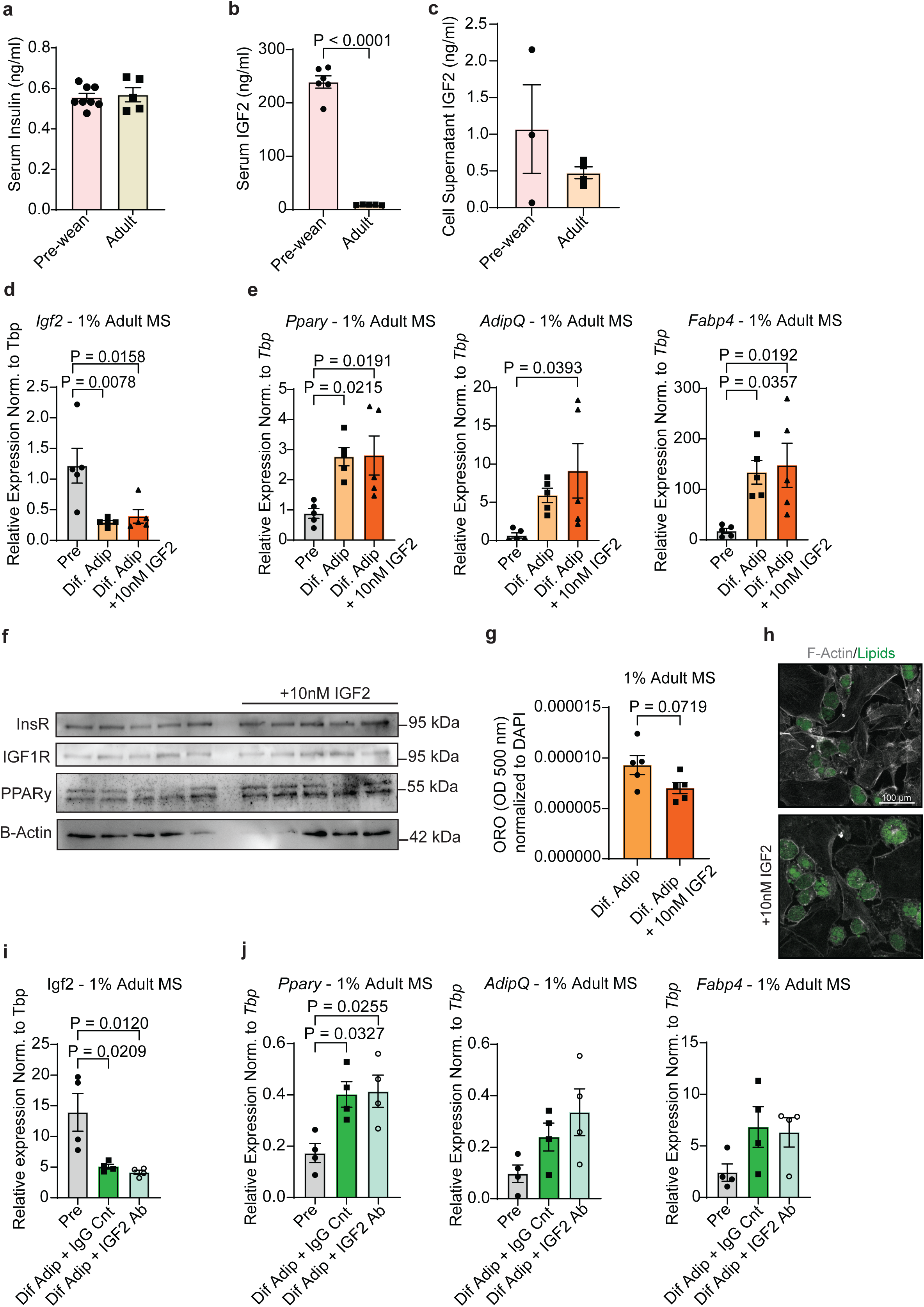
IGF2 does not alter differentiation of preadipocytes despite the substitution of FBS with mouse serum. Circulating levels of (**a**) insulin (n=8 pre-wean and n=5 adult) and (**b**) IGF2 (n=6 pre-wean and n=5 adult) in pre-wean and adult mice. (**c**) Secreted IGF2 levels in cell culture supernatant of pre-wean and adult primary scWAT preadipocytes (n= 3 pre-wean and n=4 adult). Primary adult preadipocytes were supplemented with (Dif. Adip + 10 nM IGF2) or without (Dif. Adip) 10 nM IGF2 during differentiation. mRNA expression levels of (**d**) *Igf2* (n=5) and (**e**) adipogenic markers (n=5). (**f**) Western blot of differentiated adipocytes for InsR, IGF1R, PPARγ. B-Actin was used as loading control (n=5). (**g**) Lipid quantification by Oil Red O (ORO) (n=5). (**h**) Immunocytochemistry of F-Actin (grey) and Lipids (green) (n=4). Primary pre-wean preadipocytes were treated with 1 µg/ml IgG (Dif. Adip + IgG Cnt) or IGF2 neutralizing antibody (Dif. Adip IGF2 Ab) during differentiation. MRNA expression levels of (**i**) *Igf2* (n=4) and (**j**) adipogenic markers (n=4). Data shown by mean ± SEM.

**Supplementary Figure 4:**
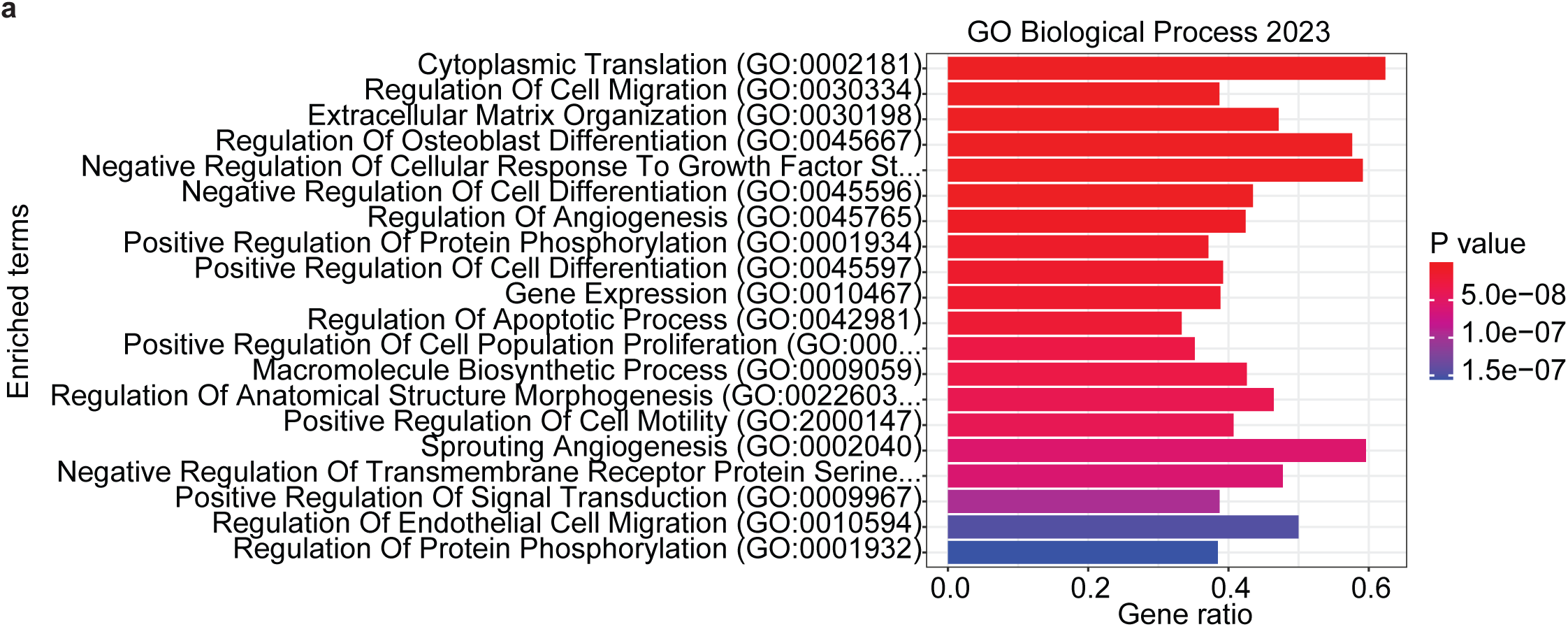
*Igf2* enriched cells express genes regulating ECM organization and cell proliferation. (**a**) Gene enrichment analysis of *Igf2* enriched vs low cell clusters from Fig. 3b for Gene Ontology (GO) Biological Process.

